# Genetic diversity and antimicrobial susceptibility pattern of Shiga toxin-producing *Escherichia coli* and *Campylobacter* spp. isolated from healthy goats in southern Thailand

**DOI:** 10.64898/2026.04.18.719346

**Authors:** Ratchakul Wiriyaprom, Ruttayaporn Ngasaman, Domechai Kaewnoi, Sakaoporn Prachantasena

## Abstract

Foodborne illness is a significant public health concern worldwide. Shiga toxin–producing *Escherichia* coli and *Campylobacter* species are recognized as important zoonotic bacterial pathogens contributing to human infections through the food chain, particularly via foods of animal origin. Although goat meat is in high demand in the southern region of Thailand, studies on foodborne pathogens in this livestock species remain limited. The current study aimed to (i) determine the antimicrobial susceptibility of *Campylobacter* spp. and STEC isolated from goats, and (ii) analyze the genetic relationships among *Campylobacter* spp. And *E. coli* O157 isolates obtained from different sources. *Campylobacter jejuni* and *C. coli* isolates were characterized based on sequences of seven housekeeping genes using the Achtman multilocus sequence typing scheme. For *E. coli* O157:H7, core genome multilocus sequence typing analysis was performed using whole-genome sequencing data. Genetic diversity was observed among *C. jejuni*, whereas a clonal population structure was detected in *C. coli* and *E. coli* O157:H7. Overlapping genetic characteristics were observed between *C. jejuni* isolates from goats and those previously reported in livestock and humans in Thailand. Among *Campylobacter* species, resistance to fluoroquinolones, including ciprofloxacin and nalidixic acid, was observed, whereas resistance to fosfomycin was most frequently detected in Shiga toxin–producing *E. coli*. Tetracycline-resistant isolates were identified in both *Campylobacter* species and Shiga toxin–producing *E. coli* at moderate levels. A multidrug-resistant pattern was observed only in *C. coli*, whereas no multidrug-resistant *C. jejuni* or Shiga toxin–producing *E. coli* isolates were detected. These findings indicate that healthy goats may serve as potential reservoirs of zoonotic pathogens and antimicrobial resistance in southern Thailand, where goat meat is frequently consumed.

## Introduction

In Thailand, goat products are generally unpopular among consumers. However, there is a high demand for goat products in the southern region of the country. According to the Department of Livestock Development, this area has the largest number of goat farmers. Most farms operate under semi-intensive or free-grazing systems. Previous studies have shown that the presence of *Campylobacter* in goats is associated with small-scale farms. This type of farming may increase the risk of exposure to environmental contaminants and pathogen transmission. Despite appearing healthy, livestock can transmit foodborne pathogens into the human food chain through direct contact, contaminated meat, or dairy products (1).

*Campylobacter* is a Gram-negative, flagellated, corkscrew-shaped zoonotic bacterium that primarily inhabits the digestive tract of certain animal species. In humans, it is recognized as a foodborne pathogen that can cause illnesses ranging from self-limiting gastroenteritis to serious post-infectious sequelae (2). Among *Campylobacter* species, *C. jejuni* and *C. coli* are most commonly associated with human illness. Human infection is usually linked to the consumption or handling of chicken meat products, exposure to contaminated objects, or contact with poultry (3). Although poultry are widely recognized as the most important food animals carrying *Campylobacter*, this organism has also been reported in other farm animals such as swine, cattle, sheep, and goats (4, 5).

Unlike *Campylobacter* spp., *Escherichia coli* can be found in a variety of environments, including the gastrointestinal tract, food, water, and vegetables (6). Most strains are harmless to humans and animals, but certain strains are considered foodborne pathogens (7). Among the pathogenic groups, Shiga toxin–producing *E. coli* (STEC) is the most virulent, causing severe disease in humans. Human illness-associated STEC is classified into several serogroups, including O26, O45, O103, O111, O121, O145, and O157, with O157 being the most well recognized because of its virulence (8). Accordingly, STEC is categorized as O157 STEC and non-O157 STEC. Hemolytic uremic syndrome and hemorrhagic colitis are among the most common severe outcomes of STEC infection (9). Outbreaks of O157 STEC infection are often associated with the consumption of beef and beef products, whereas non-O157 STEC has more frequently been reported in other types of meat such as lamb, pork, and chicken (10–12).

The emergence of antimicrobial resistance in foodborne pathogens is a major public health concern. Many studies have demonstrated that the misuse and overuse of antimicrobial agents in veterinary practice and animal husbandry accelerate the development of resistant strains. Multidrug-resistant organisms not only reduce therapeutic efficacy in human medicine but also facilitate the dissemination of resistance genes into the environment through horizontal gene transfer (13). Antimicrobial resistance in STEC and *Campylobacter* spp. isolated from small ruminants has been previously reported, with phenotypic resistance to fluoroquinolones, tetracyclines, and sulfamethoxazole observed at varying prevalence levels (4).

To understand the evolutionary dynamics and epidemiology of foodborne pathogens, the study of genetic relatedness and diversity among bacterial isolates from various sources is an effective approach. Multilocus sequence typing (MLST) has been widely used to genotype foodborne pathogens isolated from livestock by comparing sequences of seven to eight conserved housekeeping genes (14). In recent years, whole-genome sequencing (WGS) has been increasingly applied to identify clonal lineages, trace sources of bacterial origin, and detect virulence genes (15). Core genome MLST is a powerful method for analyzing genetic relatedness using WGS data. Its discriminatory power and data applicability are superior to conventional MLST because it incorporates more than a thousand core genes (16).

In Thailand, data on the genetic diversity and epidemiology of foodborne pathogens isolated from goats remain limited. The objectives of the present study were (i) to determine the antimicrobial susceptibility of *Campylobacter* spp. and STEC isolated from goats and (ii) to analyze the genetic relationships among *Campylobacter* spp. and *E. coli* O157 isolates obtained from different sources.

## Materials and methods

### Collection of bacterial strains

In total, 91 foodborne pathogen isolates, including *C. jejuni* (n = 13), *C. coli* (n = 14), non-O157 STEC (n = 54), and O157 STEC (n = 8), were investigated (S1 Table). The bacterial strains were part of *Campylobacter* and STEC surveillance in meat goats in Southern Thailand, as reported in previous studies (1, 17). The genus and species of all isolates were confirmed using molecular techniques. *Campylobacter* species were identified by the *hip*O (for *C. jejuni*) and *gly*A (for *C. coli*) genes (18). STEC was determined by the presence of Shiga toxin–encoding genes (*stx1* or *stx2*). Identification of O157 strains was defined by the presence of the *rfb* (O antigen–encoding) gene specific for *E. coli* O157 (19). All bacterial isolates were stored at −80°C prior to further analysis.

### Genetic characterization of *Campylobacter* spp

Frozen stocks of *Campylobacter* spp. and *E. coli* were recovered on blood agar (Oxoid Ltd., UK). *Campylobacter* spp. and *E. coli* isolates were incubated for 48 hours at 42°C under microaerobic conditions and for 24 hours at 37°C under aerobic conditions, respectively. DNA was extracted from fresh bacterial colonies using a DNA extraction kit (Qiagen GmbH, Germany). Nucleotide quality and quantity were assessed using a NanoDrop spectrophotometer (Thermo Fisher Scientific Inc., USA).

MLST of *C. jejuni* and *C. coli* was conducted by sequencing seven housekeeping genes, according to Jolley et al. (20). Primer sets for amplification and sequencing of internal fragments of these genes were used as described by Dingle et al. (21). Polymerase chain reaction (PCR) products were purified using a PCR extraction kit (Yeastern Biotech Co., Ltd., Taiwan) and submitted for Sanger sequencing. Allele numbers, sequence types (STs), and clonal complexes were assigned according to the MLST database (20). Phylogenetic analysis was performed using Molecular Evolutionary Genetics Analysis (MEGA) software (22).

### WGS analysis of *E. coli* serotype O157

Frozen stocks of *E. coli* O157 were recovered on trypticase soy agar (Thermo Fisher Scientific Inc., USA) and incubated for 24 hours at 37°C under aerobic conditions. A single colony from each plate was subsequently inoculated into tryptic soy broth (Merck KGaA, Germany) and incubated to increase bacterial cell numbers for downstream analyses. Bacterial suspensions were processed for DNA extraction using a commercial DNA extraction kit (Qiagen GmbH, Germany). The quality and quantity of extracted DNA were measured using a NanoDrop spectrophotometer (Thermo Fisher Scientific Inc., USA) prior to submission for WGS.

WGS of five *E. coli* O157 isolates was performed using the HiSeq 3000 platform (Illumina, CA, USA). Short sequence reads were assembled using Unicycler (version 0.5.1) prior to ST designation. Raw Illumina reads were quality-checked using FastQC (v0.11.9) and trimmed using Trimmomatic (v0.39), retaining reads of ≥50 bp with an average quality of ≥Q30. Assembly validation was conducted using Burrows–Wheeler Aligner. Achtman MLST and core genome (cgMLST) schemes of the target isolates were analyzed using EnteroBase

(23). Seven housekeeping genes were used to define STs in the Achtman MLST scheme, whereas 2,513 gene loci were used to define the cgMLST profiles.

The assembled genomes of five *E. coli* O157 isolates from the current study, together with 17 *E. coli* strains isolated from Asian countries, were analyzed using the cgMLST scheme. Whole-genome sequences from other studies were retrieved from an online database (24). A minimal spanning tree algorithm was used to illustrate genetic relationships among cgMLST profiles and visualized using GrapeTree (25). Antimicrobial resistance genes, genetic mutations, and virulence factors were also screened and characterized using the EnteroBase platform (26) and ABRicate within the Galaxy platform (27, 28).

### Antimicrobial susceptibility testing

Antimicrobial susceptibility of *Campylobacter* spp. and STEC was determined by the broth microdilution method according to Clinical and Laboratory Standards Institute (CLSI) guidelines (29). Five and eight antimicrobial agents were selected for *Campylobacter* spp. and STEC, respectively. *Campylobacter* jejuni ATCC 33560 and *E. coli* ATCC 25922 were used as quality control strains.

Antimicrobial susceptibility testing of *Campylobacter* spp. was performed against ciprofloxacin (0.015–64 µg/mL), erythromycin (0.03–64 µg/mL), gentamicin (0.12–32 µg/mL), nalidixic acid (1–256 µg/mL), and tetracycline (0.06–64 µg/mL). Minimum inhibitory concentration (MIC) breakpoints for these agents were interpreted according to the National Antimicrobial Resistance Monitoring System for Enteric Bacteria (NARMS) guidelines (30).

STEC isolates were tested against eight antimicrobial agents: amikacin (0.5–64 µg/mL), ampicillin (1–32 µg/mL), ceftriaxone (0.25–64 µg/mL), ciprofloxacin (0.015–4 µg/mL), fosfomycin (0.12–64 µg/mL), gentamicin (0.25–16 µg/mL), meropenem (0.06–16 µg/mL), and tetracycline (2–32 µg/mL). For *E. coli*, the European Committee on Antimicrobial Susceptibility Testing (EUCAST) guidelines were used to interpret MIC breakpoints for amikacin, ampicillin, ceftriaxone, ciprofloxacin, fosfomycin, gentamicin, and meropenem (31), while tetracycline breakpoints were determined according to NARMS.

## Results

### Genetic characterization of *Campylobacter* spp. and STEC serotype O157 isolated from goats

Amongst 27 *Campylobacter* spp. isolates, 19 STs were identified (Fig. 1). Seven STs were clustered into five clonal complexes, while seven STs could not be assigned to any known clonal complex (S2 Table). Novel alleles (*pgm* 1207, *gltA* 706, and *gltA* 689) and new STs (ST-11979, ST-11776, ST-11768, ST-11956) were assigned in the PubMLST database. The most common clonal complex among *C. coli* isolates was CC-828, whereas no predominant clonal complex was identified among *C. jejuni*. Close genetic relationships were observed among *Campylobacter* isolates from farms B5, F2, F3, F5, and H4. In addition, genetically related isolates were identified across different farms, including ST536 from farms F2 and H10, ST464 from farms F3 and D3, and ST860 from farms 4 and 8 (Fig. 1).

**Fig. 1.**
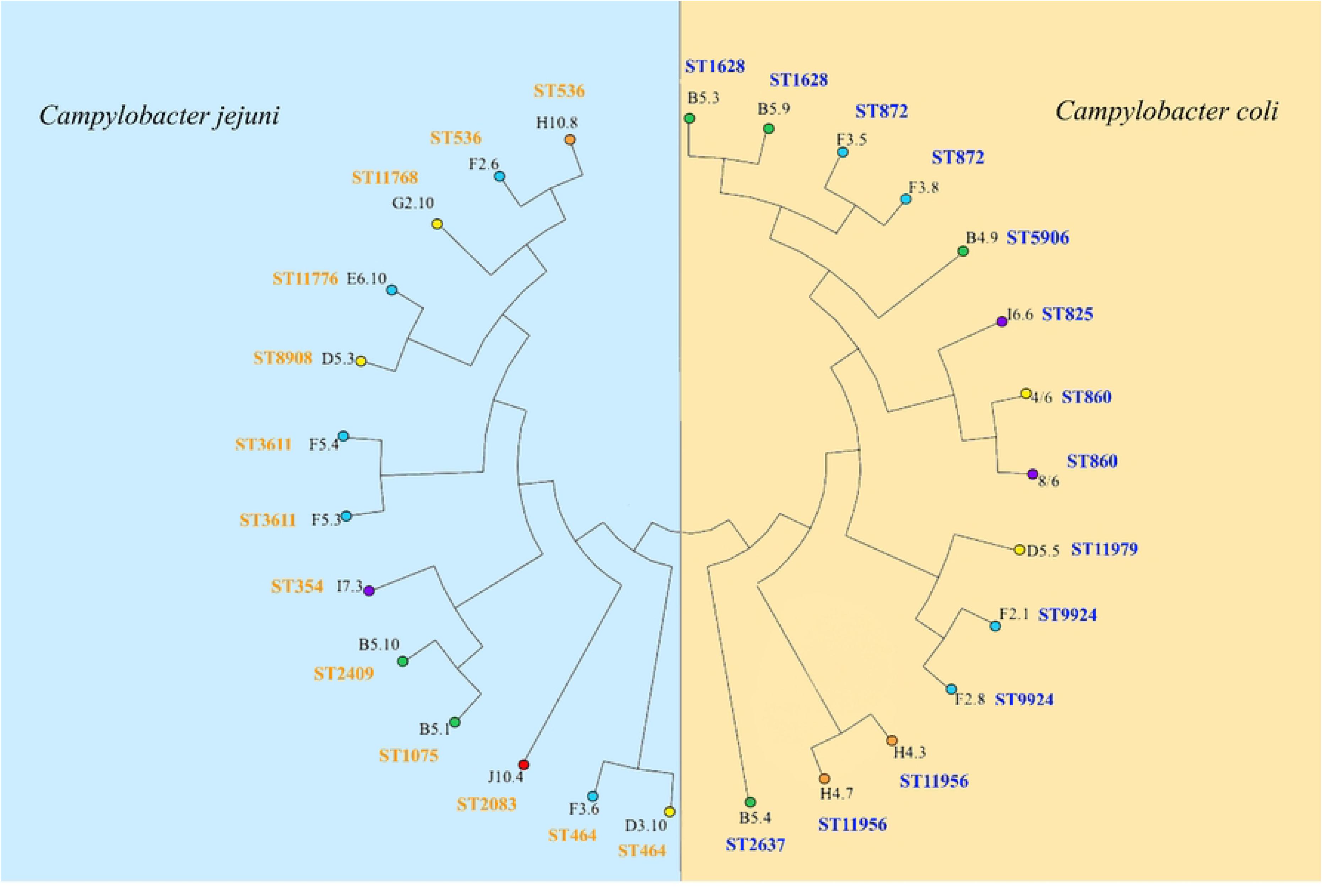
Neighbor-joining tree showing the genetic relationships of *C. jejuni* (blue background) and *C. coli* (yellow background) based on MLST data. Tip labels indicate sample name (black) and ST (blue and orange). Balloon colors at the end of each tip represent provinces of sample origin 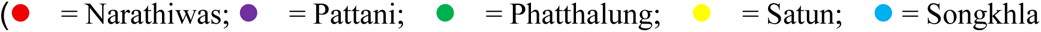; 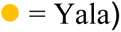.

Five *E. coli* serotype O157 isolates were selected for WGS. All isolates were confirmed as O157:H7. Under the Achtman MLST scheme, all isolates were identified as ST11, belonging to the ST11 complex, whereas three distinct STs (cgST372294, cgST372302, and cgST372320) were identified by the cgMLST scheme. Isolates originating from different farms were assigned to distinct STs, whereas isolates obtained from the same farm were classified within the same ST (S2 Table). The minimum spanning tree based on cgMLST allelic profiles of five *E. coli* O157 isolates and 17 additional *E. coli* isolates from Asian countries revealed the genetic relationships among the isolates (Fig. 2). cgST372294 and cgST372320 showed close genetic relatedness despite being isolated from geographically distant locations. By contrast, isolates assigned to cgST372302 showed greater allelic distance from cgST372294, even though both cgSTs originated from the same province (S1 Table).

**Fig. 2.**
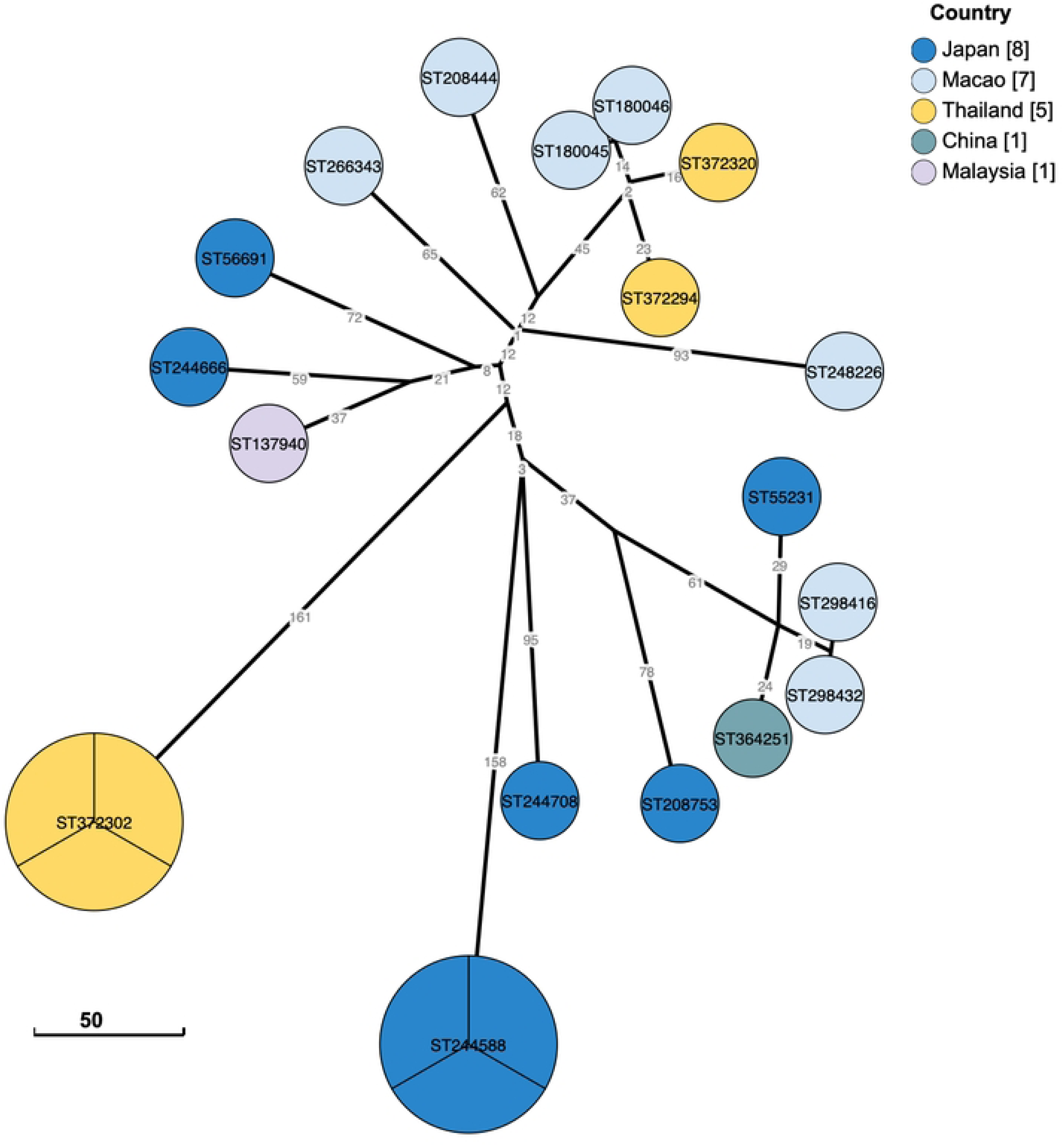
Minimum-spanning tree of five *E. coli* O157 strains from this study constructed together with *E. coli* O157 genomes reported in Asian countries and retrieved from EnteroBase. Each node represents a distinct cgMLST profile, and node size corresponds to the number of isolates. Yellow nodes indicate isolates from this study, whereas nodes in other colors indicate isolates from other sources.

### Detection of antimicrobial resistance genes and virulence factors of *E. coli* serotype O157

Whole-genome sequences were analyzed to identify antimicrobial resistance determinants and virulence characteristics of *E. coli* O157 isolates. A point mutation in the *glpT* gene (E448K), associated with fosfomycin resistance, and the multidrug transporter gene *mdfA* were detected in all five isolates. No genetic determinants responsible for resistance to aminoglycosides, macrolides, phenicols, quinolones, sulfonamides, or tetracyclines were detected. In addition, genes encoding β-lactamase inhibitors, carbapenemases, or extended-spectrum β-lactamases were not identified.

For virulence factors, the *fimH* gene in all isolates was assigned to the fimH82 type. Virulence-associated genes, including *stx2* and *eae*, were detected in all isolates. By contrast, *stx1*, *ipaH*, *plnv*, heat-stable enterotoxin (ST), and heat-labile enterotoxin (LT) genes were not detected.

### Phenotypic antimicrobial susceptibility of *Campylobacter* spp. and STEC from goats

In this study, 29 *Campylobacter* spp. and 62 STEC isolates were tested for antimicrobial susceptibility. Among *Campylobacter* spp., resistance to fluoroquinolones (i.e., ciprofloxacin and nalidixic acid) was observed at the highest proportion, followed by erythromycin and tetracycline, whereas gentamicin-resistant *Campylobacter* was rarely detected (Table 1). MIC values for *C. coli* showed a higher trend than those for *C. jejuni* (Table 2). Among STEC isolates, no resistance to ceftriaxone, meropenem, ciprofloxacin, or gentamicin was observed, whereas resistance to fosfomycin was most frequent, followed by tetracycline. Resistance to amikacin and ampicillin was observed in one isolate. Antimicrobial resistance was more prevalent in non-O157 STEC isolates than in O157 STEC isolates (Table 3). Multidrug resistance is defined as resistance to three or more classes of antimicrobial agents. A multidrug-resistant pattern was observed only in *C. coli*, whereas no multidrug-resistant *C. jejuni* or STEC isolates were detected (Table 4).

**Table 1.** Occurrence of antimicrobial-resistant *Campylobacter* spp. and STEC isolated from goats.

| Antimicrobial agents | Resistant isolates, n (%) |  |  |  |
| --- | --- | --- | --- | --- |
|  | *C. jejuni* | *C. coli* | Non-O157 STEC | O157 STEC |
| Amikacin | n/a | n/a | 1 (1.9) | 0 (0.0) |
| Ampicillin | n/a | n/a | 1 (1.9) | 0 (0.0) |
| Ceftriaxone | n/a | n/a | 0 (0.0) | 0 (0.0) |
| Fosfomycin | n/a | n/a | 6 (11.1) | 2 (25.0) |
| Meropenem | n/a | n/a | 0 (0.0) | 0 (0.0) |
| Ciprofloxacin | 12 (92.3) | 16 (100) | 0 (0.0) | 0 (0.0) |
| Gentamicin | 0 (0.0) | 1 (6.3) | 0 (0.0) | 0 (0.0) |
| Tetracycline | 3 (23.1) | 12 (75.0) | 5 (9.3) | 0 (0.0) |
| Erythromycin | 7 (53.8) | 12 (75.0) | n/a | n/a |
| Nalidixic acid | 11 (84.6) | 16 (100) | n/a | n/a |

**Table 2.**
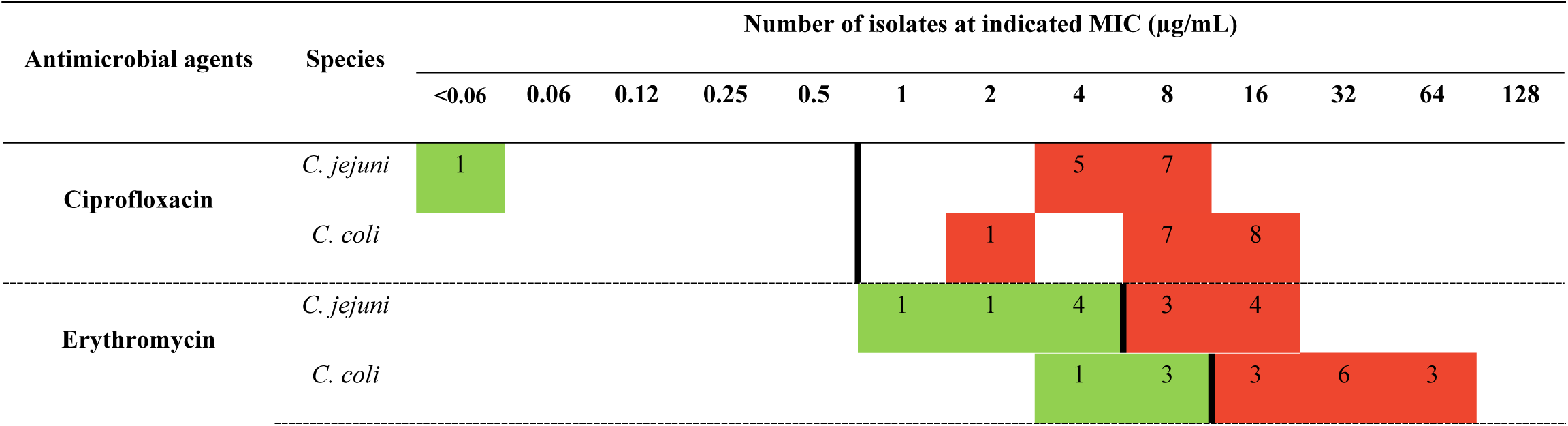
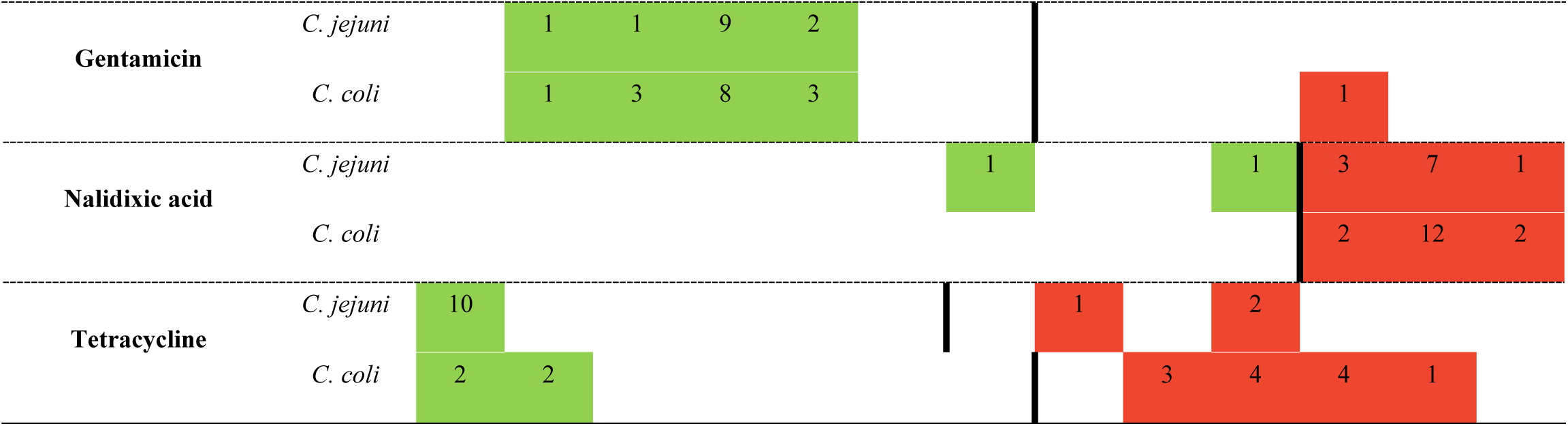
Distribution of MIC values for *C. jejuni* and *C. coli* isolated from goats. Numbers indicate the number of isolates at each MIC value. Vertical lines represent clinical breakpoints defining susceptible, intermediate, and resistant categories. Breakpoints vary by antimicrobial agent according to CLSI guidelines. Green shading indicates susceptible isolates, whereas red shading indicates resistant isolates.

**Table 3.**
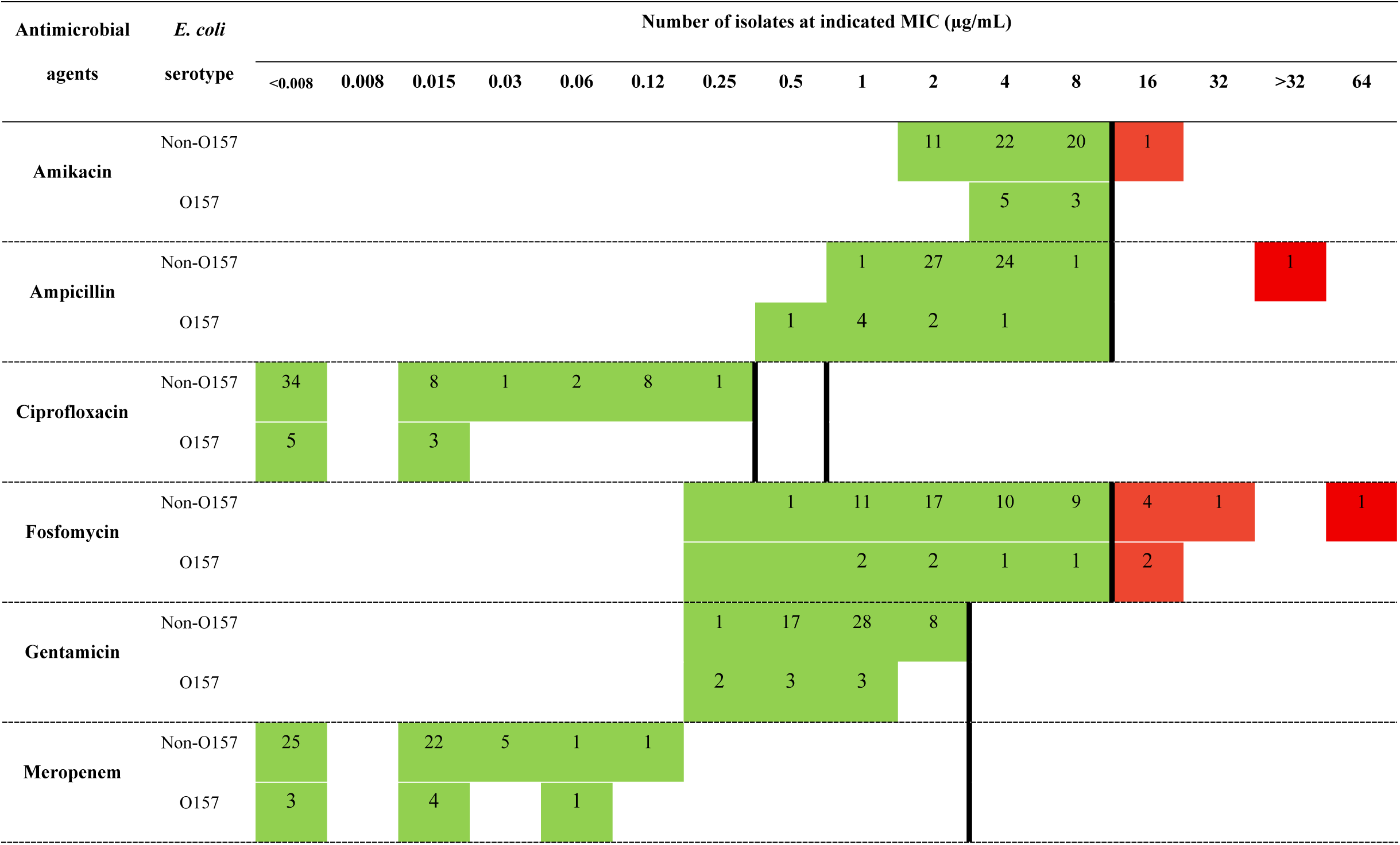
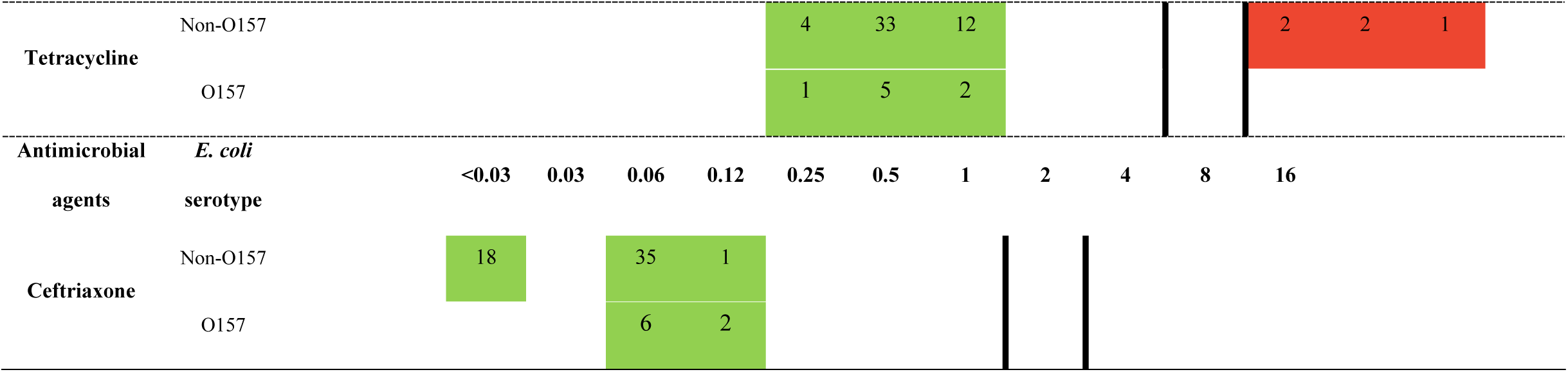
Distribution of MIC values for non-O157 and O157 STEC isolated from goats. Numbers indicate the number of isolates at each MIC value. Vertical lines represent clinical breakpoints defining susceptible, intermediate, and resistant categories. Breakpoints vary by antimicrobial agent according to CLSI guidelines. Green shading indicates susceptible isolates, whereas red shading indicates resistant isolates.

**Table 4.** Antimicrobial resistance profiles of *Campylobacter* spp. and STEC isolates. Resistance profiles represent combinations of antimicrobial agents to which isolates were resistant. Data are presented as number of isolates (percentage). Abbreviations: A, ampicillin; Ami, amikacin; C, ciprofloxacin; E, enrofloxacin; F, fosfomycin; G, gentamicin; N, nalidixic acid; T, tetracycline. “-” indicates not applicable.

| Antimicrobial resistance profiles | Bacterial species/serogroup, n (% of total isolates) |  |  |  |
| --- | --- | --- | --- | --- |
|  | *C. jejuni* | *C. coli* | Non-O157 STEC | O157 STEC |
| CEGNT | 0 (0.0) | 1 (6.3) | - | - |
| CENT | 1 (7.7) | 7 (43.8) | - | - |
| CEN | 6 (46.2) | 4 (25.0) | - | - |
| CNT | 1 (7.7) | 4 (25.0) | - | - |
| CN | 3 (23.1) | 0 (0.0) | - | - |
| C | 1 (7.7) | 0 (0.0) | 0 (0.0) | 0 (0.0) |

| Antimicrobial<br>resistance profiles | Bacterial species/serogroup, n (% of total isolates) |  |  |  |
| --- | --- | --- | --- | --- |
|  | *C. jejuni* | *C. coli* | Non-O157 STEC | O157 STEC |
| T | 0 (0.0) | 0 (0.0) | 3 (5.6) | 0 (0.0) |
| F | - | - | 5 (9.3) | 2 (25.0) |
| Ami | - | - | 1 (1.9) | 0 (0.0) |
| TA | - | - | 1 (1.9) | 0 (0.0) |
| TF | - | - | 1 (1.9) | 0 (0.0) |
| Susceptible | 1 (7.7) | 0 (0.0) | 43 (79.6) | 6 (75.0) |
| **<u>Total</u>** | **<u>13</u>** | **<u>16</u>** | **<u>54</u>** | **<u>8</u>** |

## Discussion

This study investigated the genotypic and phenotypic characteristics of *Campylobacter* spp. and STEC isolated from healthy goats in the lower part of southern Thailand. Genotypic characterization of *Campylobacter* spp. and O157 STEC was conducted using the Achtman MLST and cgMLST schemes, respectively. A total of 91 bacterial isolates, including *C. jejuni*, *C. coli*, non-O157 STEC, and O157 STEC, were subjected to antimicrobial susceptibility testing.

### Genetic diversity of *Campylobacter* spp. isolated from goat

MLST revealed considerable genetic diversity among *C. jejuni* isolates obtained from goats across different geographic locations. Seven STs were assigned to five clonal complexes and one novel ST, indicating a complex population structure. However, several isolates from different farms shared identical STs, including F2.6 and H10.8 (ST536) and D3.10 and F3.6 (ST464), suggesting possible dissemination of specific strains across farms. Among *C. coli* isolates, CC-828 was the predominant clonal complex, reflecting a more clonal population structure. Previous studies in Thailand have reported several STs identified in the present study in poultry and other livestock species, indicating overlapping reservoirs and potential shared transmission routes among domestic animals. Phu et al. (32) described MLST profiles of *Campylobacter* spp. from commercial and native chickens in the upper part of southern Thailand, which partially overlapped with the goat-derived isolates identified here. Similarly, CC-353, previously reported as predominant in the broiler production chain in central and eastern Thailand, was also detected among these isolates (33). Importantly, these findings highlight the zoonotic potential of goat-derived *Campylobacter* spp. in Thailand. Several STs identified in this study (i.e., ST-860, ST-464, ST-1075, ST-2409, ST-8908, ST-354, and ST-2083) have also been reported in human clinical isolates in Thailand (20), supporting the role of goats as a potential reservoir of *Campylobacter* spp.

### Genetic characterization of *E. coli* O157

In addition to the STEC isolates recovered from goats, five *E. coli* O157:H7 isolates were further analyzed using WGS. Clonality among the five isolates was determined using the Achtman MLST scheme, which classified all isolates as ST11. ST11 has been described as the predominant ST associated with *E. coli* O157:H7 and has been widely reported among virulent STEC strains in severe human infections (34). A high degree of genetic similarity was observed between goat-derived *E. coli* O157:H7 isolates and food-derived strains reported in previous studies (S3 Table). Comparative cgMLST analysis with *E. coli* O157:H7 strains from the Asian region revealed that a strain from Macao exhibited the closest genotypic relationship to the isolates identified here, whereas strains from China and Japan appeared more distantly related. Similar genomic relatedness among *E. coli* O157:H7 isolates from different sources has been reported in previous studies, highlighting the potential role of food-producing animals as a source of this pathogen (35, 36).

All five isolates carried the *stx2* and *eae* genes, which are key virulence determinants of STEC. STEC strains are typically identified by the presence of *stx1* or *stx2* genes through molecular methods. Expression of these genes results in the production of Stx1 and Stx2 toxins, respectively. *Escherichia coli* strains producing Stx2 have been strongly associated with severe human illnesses, including watery or bloody diarrhea, hemorrhagic colitis, and hemolytic uremic syndrome, whereas strains producing Stx1 alone are generally linked to milder clinical symptoms (37). In addition, the *eae* gene encodes an adhesion protein that mediates intimate attachment to host epithelial cells and plays an important role in bacterial pathogenicity (38). These findings indicate that goats could represent an important reservoir of *E. coli* O157:H7. The presence of virulent STEC strains in goats highlights the potential risk of zoonotic transmission.

### Phenotypic antimicrobial susceptibility patterns of *Campylobacter* spp. and STEC

Our findings revealed high levels of resistance to fluoroquinolones, macrolides, and tetracyclines in *C. coli*, whereas resistance rates were slightly lower in *C. jejuni*. In addition, very low levels of gentamicin resistance were observed in both *Campylobacter* species. Increasing fluoroquinolone resistance in *Campylobacter* has previously been documented in Thailand. For instance, Boonmar et al. (39) reported low levels of fluoroquinolone-resistant *C. jejuni* from human and poultry sources. High levels of macrolide resistance in *C. jejuni* from human and poultry sources have also been reported. Padungtod et al. (40) similarly described relatively high fluoroquinolone resistance in *C. jejuni* and *C. coli* isolated from humans, animals, and animal products in northern Thailand. Previous studies have indicated that erythromycin-resistant *Campylobacter* strains are more prevalent in the swine production chain and in humans, whereas lower erythromycin resistance has been reported in cattle and poultry production systems (40). In this study, the most prevalent antimicrobial resistance pattern in *C. jejuni* was ciprofloxacin–erythromycin–nalidixic acid, whereas ciprofloxacin–erythromycin–nalidixic acid–tetracycline was the most common pattern in *C. coli*. These findings differ slightly from a previous study conducted in southern Thailand (41), which reported ciprofloxacin–nalidixic acid–tetracycline as the predominant resistance pattern in *C. jejuni* and ciprofloxacin–erythromycin–nalidixic acid–tetracycline in *C. coli* isolated from commercial and native chickens (41). In this study, multidrug-resistant *Campylobacter* was also detected among *C. coli* isolates, although at a relatively low level. Previous studies have reported a higher proportion of multidrug-resistant *Campylobacter* among isolates from pigs, whereas lower levels have been observed in cattle (40).

Low levels of antimicrobial resistance were observed among STEC isolates in this study. However, non-O157 STEC exhibited higher resistance rates than O157 STEC isolates. Overall, fosfomycin resistance was most frequently observed, followed by tetracycline. These findings are consistent with previous studies in Japan (42), which reported relatively low levels of antimicrobial resistance in O157 STEC compared with non-O157 STEC. Similarly, higher resistance rates in non-O157 STEC have been reported in isolates from humans and livestock in China (43). In addition, studies from Japan have indicated that multidrug resistance patterns are more commonly associated with non-O157 STEC than with O157 STEC (42). O157:H7 STEC isolates obtained from cattle and cattle products in the United States have been reported to be susceptible to several antimicrobial agents, including amikacin, ceftriaxone, ciprofloxacin, fosfomycin, streptomycin, and tetracycline (44). By contrast, a study conducted in China showed that tetracycline resistance is most prevalent among non-O157 STEC isolates (43). Interestingly, all isolates in a study from Brazil were resistant to tetracycline, which is substantially higher than the rate observed here (45). No multidrug-resistant STEC isolates were identified in this study. By contrast, a study from Egypt reported that 57.4% of O157 STEC isolates were multidrug-resistant, whereas another study in China found multidrug resistance in 28.5% of non-O157 STEC isolates (43, 46). Furthermore, 65.6% of non-O157 STEC isolates from Brazil were resistant to three or more classes of antimicrobials (45).

Whole genome sequence analysis of O157 STEC identified a point mutation in the *glpT* gene, which encodes a glycerol-3-phosphate transport system associated with fosfomycin uptake (47). However, phenotypic resistance to fosfomycin was observed in only two of eight O157 STEC isolates. Mutations in the *glpT* gene associated with phenotypic fosfomycin resistance in *E. coli* have been previously described (8). Alternative fosfomycin transport systems, including expression of the *uhpT* gene, may compensate for *glpT* function (49). The detection of fosfomycin resistance at both phenotypic and genotypic levels highlights a potential public health concern because fosfomycin is considered a drug of choice for the treatment of multidrug-resistant *E. coli* infections and pediatric STEC infections (50).

Genetic characterization of *Campylobacter* spp. and STEC demonstrated epidemiological relationships among these foodborne pathogens in goats, which may serve as a reservoir for human infection. This study highlights the role of healthy goats as potential reservoirs of zoonotic pathogens and antimicrobial resistance. The occurrence of antimicrobial-resistant *Campylobacter* in food-producing animals may indicate a risk to public health through transmission via the food chain, emphasizing the importance of antimicrobial resistance surveillance within a One Health framework.

## Acknowledgments

We thank all goat farmers for providing samples for this study. In addition, we would like to thank Dr. Pharanai Sukhumungoon for the guidance on laboratory procedures.

